# Historic Overexploitation, Genetic Erosion and Local extinction: Palaeogenomic Insights into the Decline of *Eubalaena glacialis* in the Northeast Atlantic

**DOI:** 10.1101/2025.11.24.690181

**Authors:** Gonçalo Espregueira Themudo, Alba Rey-Iglesia, Inês M. dos Santos, Rebecca L.S.T. Netels, Alfredo López, Jose Matínez-Cedeira, Rute R. da Fonseca, Paula F. Campos

## Abstract

The North Atlantic right whale (*Eubalaena glacialis*) was the primary target for early industrial whalers, particularly the Basques. Once widespread across temperate and subpolar waters on both sides of the North Atlantic, the species is now functionally extinct in the Northeast Atlantic and listed as critically endangered by the IUCN. Despite their fragmentary remains, bones are abundant in the archaeological and historical records and reveal a long history and extensive tradition connected to the exploitation of these marine resources. Using state-of-the-art ancient DNA techniques, here we aimed at genetically characterizing the lost Northeast Atlantic population. We sequenced 17 genomes from historical specimens collected in the Cantabrian Sea, dating from the 13^th^-18^th^ century, and compared them with available genomes for 12 individuals from the surviving Northwest Atlantic population. Our analyses reveal a historically panmictic population and document a dramatic decline in genetic diversity from the Middle Ages to the present. These findings illuminate the profound impact of centuries of whaling, which eradicated the Northeast Atlantic population and severely depleted genetic diversity in the remnant population.

## Introduction

Whaling has been part of human history for millennia. It evolved from being a subsistence activity for indigenous people to a prolific commercial operation. Several coastal societies like the Vikings, Inuits and Japanese have been engaging in whale hunting for centuries, using nearly every part of the animal which played a key economic, mythological and sociological role in these societies (e.g. McCartney, 1980; McCartney and Savelle, 1993; Seersholm et al., 2016; Skovrind et al., 2024). With the advent of better tools and more advanced whaling techniques, Basque whalers established the first commercial whaling industry, with the earliest whaling dates of 1059 in Bayonne in the Gulf of Biscay and 1150 in Navarre, Northern Spain (Aguilar, 1986). The Basques pioneered a technique that required them to stay physically tethered to the whale. They used exceptionally long lines, allowing the harpooned whale to flee or dive without putting the boat at risk. When the whales were exhausted, whalers approached the animal and delivered the fatal blow. This highly effective method laid the groundwork for European and later American whaling industry (Reeves and Smith, 2007). Whale exploitation has emerged independently across various regions worldwide and by the 15^th^ century it had evolved into a significant international industry, depleting whale populations, and driving many populations and species to near extinction.

The North Atlantic right whale (NARW, *Eubalaena glacialis*), a baleen whale, which belongs to the family *Balaenidae*, along with the bowhead whale (*Balaena mysticetus*), was the main target of industrial whaling in the Cantabrian Sea (northern coast of Spain and the southwest side of the Atlantic coast of France, Figure1) during Medieval times (Rey-Iglesia et al., 2018). It was particularly targeted due to its high quality oil, but their meat was also used as a food source and the baleens had multiple utilities from the use as fishing gear to implements to clothing (Solazzo et al., 2017). Their non-aggressiveness and their seasonal preference for coastal waters, especially during breeding season, allowed boats to easily approach them.

Southern right whales (*Eubalaena australis*) show the same behaviour using Península Valdés in Argentina’s Patagonian coast as breeding ground (Sueyro et al., 2018). In addition, their high blubber content meant that corpses floated and made them a preferred target for whalers. The latter characteristic possibly gave rise to its current common name, right whale, as it was the "right" whale for whalers to hunt.

This species was once distributed all over the sub polar and temperate waters of the North Atlantic (Figure 1). However today, they are functionally extinct on the Northeast Atlantic coast and less than 400 individuals inhabit the coastal waters off the eastern United States (Pettis et al., 2022), migrating between calving grounds from Cape Fear, North Carolina, to below Cape Canaveral, Florida and feeding grounds in the Gulf of Maine and the Bay of Fundy off the coast of New England. The small effective population size and the low genetic diversity of this population when compared to Southern Right whales and other big whales have often been connected to whaling (e.g., Crossman et al., 2024; Waldick et al., 2002).

**Figure 1.**
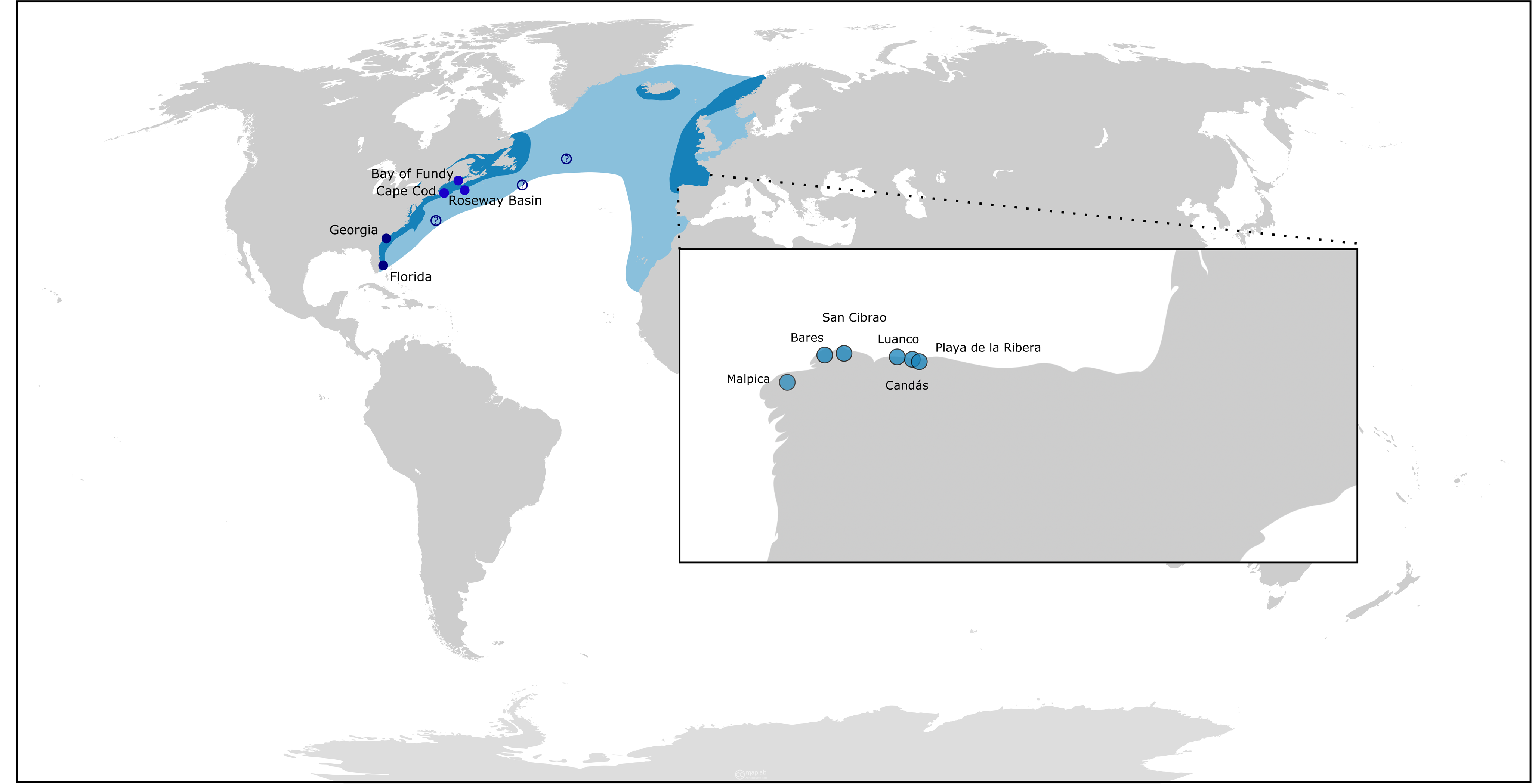
Historical distribution of the North Atlantic right whale and sampling locations. The map shows the historical range of the North Atlantic right whale (blue) and the geographical locations of historical sampling sites in Galicia and Asturias (circles), along the western Cantabrian Sea. The locations of modern samples included in this study are also indicated (squares).

NARW genetic diversity is now among the lowest recorded for any wildlife species, comparable to the cheetah and Iberian lynx (Crossman et al., 2024). Even though both NARW and Southern right whales were impacted by whaling, the northern species had a harder hit, with the population reaching the lowest ever effective population size of under 200 individuals. Analysing the demographic history of a population can provide important context for interpreting its current genetic diversity. Populations that have experienced bottleneck events typically undergo substantial reductions in genetic variation due to the combined effect of genetic drift and inbreeding (Frankham et al., 2010), and these reductions often persist even after demographic recovery, which is not the case for our study species. These demographic contractions leave long-lasting genetic signatures that will influence adaptive potential and conservation outcome of these species.

Whale bones are abundantly present in the archaeological record but its fragmentary nature hinders species or even genus identification, making tracking of species distributions and investigating changes in exploitation through time almost impossible, making them pretty much invisible in such records (Speller et al., 2016). Archaeogenomic research into the history of whaling and marine mammal hunting is vital not only for tracing the long-term exploitation of these key resources, but also for establishing essential ecological baselines of whale populations before the onset of industrial overhunting, that can then be used in conservation and management processes. This is especially important as written records from the Medieval Basque whaling period are very incomplete.

In a recent study, Béland and colleagues (2020) demonstrated that commercial whaling had a surprisingly limited impact on the overall mitochondrial and nuclear genetic diversity of Northeast Pacific humpback (*Megaptera novaeangliae*) and grey whale (*Eschrichtius robustus*) populations, despite having identified bottlenecks as direct effect of commercial whaling.

Whales play a crucial role in maintaining ecosystem balance through the "whale pump," a process in which they cycle nutrients between deep and surface waters (Roman and McCarthy, 2010). Even though most baleen whales do not actively hunt large prey, as they are filter feeders, who primarily feed on zooplankton, especially copepods and krill, they significantly influence marine food webs through their feeding habits (Freitas et al., 2025), helping to regulate prey populations and support ocean productivity. So, understanding their dynamics over time is crucial for maintaining a healthy ecosystem.

In a previous study we have sampled 70 Medieval whale specimens from off the coasts of Galizia and Asturias (Northeast Atlantic, where the NARW are now extinct) and 68 were identified as NARW using molecular tools (Rey-Iglesia et al., 2018). Here, we did whole genome re-sequencing of 39 of those samples (selected based on percentage of endogenous DNA content) aiming at characterizing the population dynamics of the NARW as well as assessing the impact of whaling activities on the genetic diversity of the species.

## Methods

### (a) Sample collection

Whale bones were collected by CEMMA (Coordinadora para o Estudo dos Mamíferos Marinos, Galicia) and collaborators during the spring and summer of 2014. Permissions were obtained from all museums and institutions to access the collections and all samples were on loan for scientific purposes. Click or tap here to enter text.Samples were collected from local museums, private collections and in underwater sampling campaigns as described in Rey-Iglesia et al (2018). Underwater sampling targeted areas described in several historical records as medieval whaling ports and were concentrated in Galicia (Bares and San Cibrao; Figure. 1). A number of written records confirm the ports of Bares, Luanco and San Cibrao as old whaling ports active between the 13th and 18th century, which allow us to confidently place our samples in this time frame (Ciriquiain-Gaiztarro, 1961; López, 2014; Valdés Hansen, 2010). All samples were stored at -20°C at CEMMA headquarters in Nigrán, Galicia, Spain, for optimal DNA preservation.

### (b) DNA extraction and library construction

All DNA extractions (Table 1) and double strand DNA Illumina library preparation steps were carried out in dedicated ancient DNA facilities at either the Natural History Museum of Denmark or at the University of Tartu in Estonia, designed for dealing with potentially degraded samples such as these, which are particularly susceptible to contamination from exogenous sources of DNA. Contamination was monitored during the extraction and PCR processes by blank controls. Total cellular DNA from 70 bone samples, was extracted from specimens previously identified as NARW according to the following protocol: 100mg of bone were powdered and incubated overnight at 37 °C in 1.0 mL of 0.5 M EDTA and 25 mg/ml proteinase K. To pellet the non-digested powder, the solution was centrifuged at 12,000 rpm for 5 min. The liquid fraction was then transferred into a Centricon micro-concentrator (30-kDa cut-off), and spun at 4000 rpm for 10 min. When the liquid was concentrated down to 200–250 μl, the DNA was purified using the MinElute PCR purification kit (Qiagen), with the following modifications: a) 13× PB buffer (Qiagen) was used for the DNA binding step; b) spins were done at 8,000 rpm with the exception of the final one at 13,000 rpm; c) in the elution step, spin columns were incubated in 25 μl EB buffer at 37 °C for 10 min, spun down, and repeated once more. The eluates from both rounds of elution were pooled.

**Table 1.** Sample information for both Medieval individuals sequenced in this study and publicly available data for extant populations. Bold indicates samples that were sexed morphologically. For * means only mitogenome data is available for these specimens.

| Sample | Locality | origin | Sex | PCA | Age |
| --- | --- | --- | --- | --- | --- |
| 1 | Malpica | this study | F | y | post 1600 |
| 2 | Bares | this study | F | y | historical |
| 3 | Luanco | this study | M | y | historical |
| 4 | Luanco | this study | M | y | historical |
| 5 | Luanco | this study | M | y | historical |
| 6 | Playa de la Ribera | this study | F | y | historical |
| 7 | Playa de la Ribera | this study | F | y | historical |
| 8 | Luanco | this study | F | y | historical |
| 9 | Luanco | this study | F | y | historical |
| 10 | Luanco | this study | M | y | historical |
| 11 | Luanco | this study | M | y | historical |
| 12 | Candás | this study | F | y | 100BCE-1800CE |
| 13 | San Cibrao | this study | F | y | 100BCE-1800CE |
| 14 | San Cibrao | this study | F | y | 100BCE-1800CE |
| 15 | San Cibrao | this study | M | y | 100BCE-1800CE |
| 16 | San Cibrao | this study | M | y | 100BCE-1800CE |
| 17 | San Cibrao | this study | M |  | 100BCE-1800CE |
| 18 | Western N Atlantic | OP205189 |  |  | modern |
| 19 | Atlantic | OP205190 |  |  | modern |
| 19 | North Atlantic | OP205191 |  |  | modern |
| 20 | Florida | OP205188 | **F** |  | modern |
| 21 | Georgia | OP205187 | **F** |  | modern |
| 22 | Georgia | OP205186 | **M** |  | modern |
| 23 | Georgia | OP205185 | **F** |  | modern |
| 24 | Georgia | OP205184 | **F** |  | modern |
| 25 | Georgia | OP205183 | **F** |  | modern |
| 26 | Georgia | OP205182 | **M** |  | modern |
| 27 | Georgia | OP205181 | **M** |  | modern |
| 28 | North Carolina | MF459656 |  |  | modern |
| 29 | Bay of Fundy | SRR22863755 | F | y | modern |
| 30 | Cape Cod | SRR22863740 | F | y | modern |
| 31 | Roseway Basin | SRR22863753 | M | y | modern |
| 32 | Bay of Fundy | SRR22863739 | F | y | modern |
| 33 | Bay of Fundy | SRR22863754 | F | y | modern |
| 34 | Bay of Fundy | SRR22863752 | M | y | modern |
| 35 | Bay of Fundy | SRR22863738 | F | y | modern |
| 36 | Bay of Fundy | SRR22863743 | F | y | modern |
| 37 | Bay of Fundy | SRR22863737 | F | y | modern |
| 38 | Florida | SRR22863736 | F | y | modern |
| 39 | Florida | SRR22863735 | F | y | modern |
| 40 | Florida | SRR22863734 | F | y | modern |

### (c) Library building and amplification

Double stranded and indexed DNA Illumina libraries were built from 39 of the extracts. A blunt-end library was constructed on 21.25 μl of DNA extract following the protocols outlined by Maricic et al. (2010) and Meyer and Kircher (2010). The amplified libraries were run on an Agilent 2100 Bioanalyzer High Sensitivity DNA chip. Library construction and sequencing was done at University of Tartu and funded by EASI Genomics (PID14751). A preliminary screening of the 39 samples was conducted, and the 17 with the highest endogenous DNA content were selected for deep sequencing.

### (d) Data processing, mapping and annotation

Base calling was performed using the Illumina software CASAVA 1.8.2, with the requirement of a 100% match to the 6-nucleotide index used during library preparation.

Raw reads were bioinformatically processed using nf-core/eager v.2.4.7 (Fellows Yates et al., 2021), a NextFlow pipeline for ancient DNA analysis. Adapter sequences were trimmed and filtered for N’s, and reads shorter than 30 bp were removed using AdapterRemoval v2.3.2 (Schubert et al., 2016). Trimmed reads as well as 12 modern specimens stored at SRA (SRR22863755, SRR22863740, SRR22863753, SRR22863739, SRR22863754, SRR22863752, SRR22863738, SRR22863743, SRR22863737, SRR22863736, SRR22863735, SRR2286373) were mapped to *Eubalaena glacialis* genome assembly (NCBI accession number GCF_028564815.1; mEubGla1) using bwa aln v0.7.17-r1188 (Li and Durbin, 2009). A consensus mitogenome sequence was generated using a minimum sequence depth of 2X and a majority rule for base calling.

Beagle files with the nuclear genome positions of single nucleotide polymorphisms (SNPs) were produced by ANGSD (Korneliussen et al., 2014) using the following options: angsd - bam $bamList -ref $REF -out ${out}.snp -C 50 -baq 2 -remove_bads 1 -uniqueOnly 1 - doCounts 1 -doGlf 2 -GL 1 -doMaf 2 -SNP_pval 1e-6 -doMajorMinor 1 -minQ 20 -minMapQ 30 -minInd 62 -minMaf 0.05 -rmTrans 1. Transitions were excluded.

### (e) Authentication and alignment

To authenticate the historical sequences obtained as genuine, we ran damageprofiler (Neukamm et al., 2021) in order to detect DNA post-mortem damage patterns typical of ancient or degraded DNA. The program uses misincorporation patterns, particularly deamination of cytosine into uracils, within a Bayesian framework. An elevated C to T substitution rate towards sequencing starts (and complementary G to A rate towards the end) is considered indicative of genuine ancient or degraded DNA.

### (f) Phylogenetic analyses

Phylogenetic relationships among these historical samples and Northwest Atlantic Right whale mitogenome sequences either retrieved from GenBank (MF459656; OP205181-OP205191) or reconstructed from SRA genomic data (see Table 1) using the same pipeline as for the ancient samples was reconstructed with Beast v2.7.6 (Bouckaert et al., 2019), using the GTR sites substitution model and a strict molecular clock. The MCMC chain ran for 10 million iterations, and the log was checked to see if the number of effective sample sizes reached at least 200 for most parameters. A median joining network (Bandelt et al., 1999)of all *E. glacialis* mitogenomes, excluding the control region, was drawn using PopArt (Leigh and Bryant, 2015). A second median joining network was reconstructed using partial dloop sequences (279bp) from all previously mentioned modern and Medieval samples, together with sequences from Iceland, Scotland and the Faroe Islands (Frasier et al., 2022), as well as a single historical specimen from Labrador, Canada. These sequences represent the 13 haplotypes (A-N) previously described (Frasier et al., 2022; Malik et al., 2000; McLeod et al., 2010; Rastogi et al., 2004).

### (g) Population structure analyses

Principal component analyses (PCAs) where obtained with PCAngsd version 1.10 (Meisner and Albrechtsen, 2018) on 17 subsets of the data, each comprising all modern samples (n=12) and one historical specimen, not allowing missing data in the latter. The number of SNPs included in each PCA is provided in table 3.

A Procrustes transformation was performed to find the best fit between the individual PCAs based on the common samples. The analysis was conducted in R using the vegan package, which provides the Procrustes() function for optimal superimposition of matrices by minimizing the sum of squared differences. Prior to transformation, data were imported and processed using the readr and dplyr packages for efficient data handling. Visualization of the Procrustes results, including the superimposed configurations and residuals, was generated using ggplot2 to facilitate interpretation of congruence between datasets.

### (h) Demographic reconstruction

We reconstructed female mitogenomic demographic histories using the Bayesian Skyline Plot (BSP) (Drummond et al., 2005). BSP analyses were performed in BEAST version 2.7.6 (Bouckaert et al., 2019) using a coalescent Bayesian Skyline and strict molecular clock with a substitution rate of 7 × 10^−9^ substitutions/site/y (Sasaki et al., 2005) and HKY substitution model with gamma distribution. The chain length was established at 1 × 10^8^ iterations, with samples drawn every 10,000 Markov chain Monte Carlo steps. After a discarded burn-in of 25%, the BSP was reconstructed and visualized in a plot with Tracer version 1.7.2 (Rambaut et al., 2018).

### (i) Genetic sexing

We determined the genetic sex of the right whale samples by calculating the ratio between depth of coverage of the X chromosome (NC_083736) and depth of coverage of the similarly sized chromosome 7 (NC_083722). We considered that a ratio above 0.8 indicated that the individual was a female, a ratio below 0.6 a male, and between the two values was undetermined.

## Results and discussion

### Data processing, mapping, annotation

The North Atlantic right whale samples were mapped to the *Eubalaena glacialis* genome assembly (NCBI accession number GCF_028564815.1; mEubGla1). An initial screening of the 39 samples identified as North Atlantic right whale (NARW) revealed substantial variation in endogenous DNA content, ranging from 0.12% to 42.13%. From these, 17 samples exhibiting the highest endogenous DNA content (2.19–37.28%) were selected for deep sequencing (Table S1). Sequencing produced a total of 1,042,078,041 reads (min 19,218,490 – max 88,904,651/ reads per sample), of which 50,275,456 mapped to the nuclear genome, while 123,599 reads mapped to the mitochondrial genome (min 1,565 – max 22,075, see Table S1). This resulted in an average depth of coverage ranging from 0.02 - 0.18x for the nuclear genome and 4.1 – 68.3x for the mitochondrial genome (Table S1).

### Authentication

Misincorporation plots (supplementary material Figure 1) show a typical pattern of higher C to T and G to A transitions a the 5’ and 3’ ends, respectively, indicating the retrieval of genuine endogenous DNA from these historical samples.

### Mitogenome phylogeny

Phylogenetic analysis of complete mitochondrial genomes (mtDNA) from 17 ancient individuals from the northeastern Atlantic (this study) and 24 modern individuals from the northwestern Atlantic (n = 41) reveals partial overlap in genetic diversity between the two groups. Several clades include both ancient and modern individuals, while others contain only ancient or only modern samples (Figure 2A). Likewise, in the median-joining network of the complete mitogenome, the most common haplotype is shared across both temporally and geographically separated populations, whereas most haplotypes are unique to either the ancient or modern groups (Figure 2B). There is no distinct separation between northeastern and northwestern haplotypes; many differ by only one or two mutations, suggesting historical gene flow and frequent admixture between the two regions. However, a marked loss of mitochondrial diversity is evident in the modern northwestern Atlantic population when compared to our Medieval Cantabrian samples, consistent with previous findings by Frasier et al. (2022), who reported genetic erosion in archaeological samples from Iceland, the Faroe Islands, and Scotland based on a small fragment of the d-loop. This phenomenon is probably due to excessive whaling during the last centuries which depleted genetic diversity in the Northwestern population and led to the extinction of the northeastern stock. The median joining network from the partial dloop sequences which include both historical Cantabrian and Northern European samples (Iceland, the Faroe Islands and Scotland) along with the 13 previously described haplotypes show a similar pattern. The analyses of this new set of 17 historical samples allowed the description of five new haplotypes in these Medieval populations (O-S).

**Figure 2.**
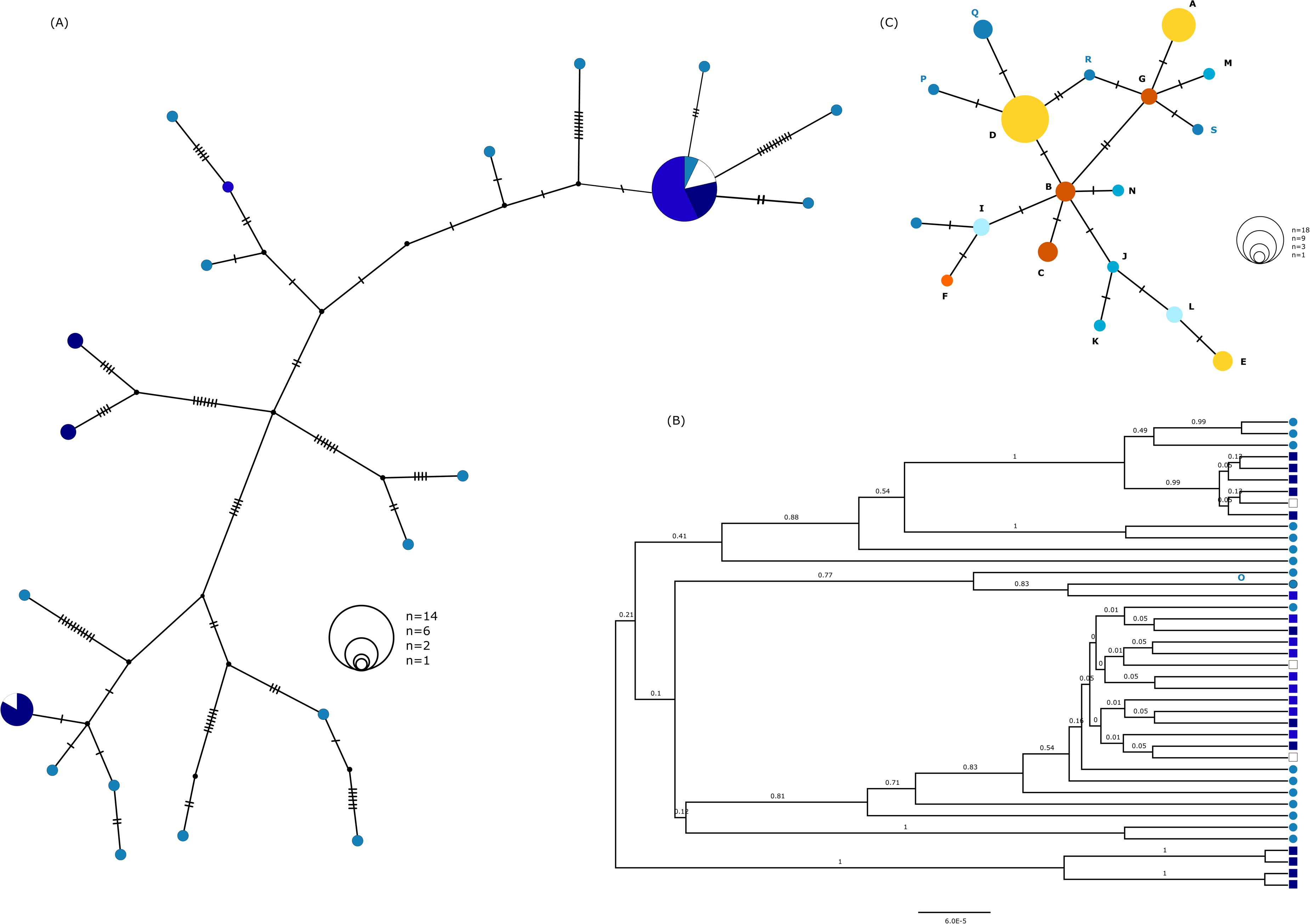
Phylogenetic relationships between historical and modern North Atlantic right whales based on complete mitogenomes and partial dloop.; (A) median-joining network generated from 41 NARW mitogenomes, comprising 17 historical and 24 modern individuals. Bayesian phylogenetic tree in which rectangles denote modern individuals and circles denote Medieval specimens. In both panels, white indicates individuals of North Atlantic origin with no precise locality, teal represents Medieval Cantabrian samples, bright blue corresponds to modern individuals from feeding grounds, and navy blue to modern individuals from breeding grounds; (C) median-joining network generated from a fragment of the dloop from 54 specimens including all 13 previously described haplogroups A-N, from all previously mentioned modern and Medieval samples, together with historical individuals from Iceland, Scotland, the Faroe Islands and a single historical specimen from Labrador, Canada. Medieval Cantabrian samples are represented in teal (haplotypes O-S); historical samples from Iceland, Faroe Islands and Scotland in blue (J,K,M,N); historical Northeast Atlantic specimens in light blue (haplotypes I,L); Northwest Atlantic samples in dark orange (haplotypes B,C,G); historical sample from Labrador in bright orange (haplotype F) and in yellow (haplotypes A, D, E) modern samples from the Northwest Atlantic and Medieval samples from the Cantabrian Sea.

### Demographic analyses

Bayesian Skyline Plot (BSP) analysis based on complete mitochondrial genomes (n = 41) revealed a pronounced and sustained decline in effective female population size (*Nₑf*) beginning approximately 5,000 years ago. The BSP curve shows a relatively stable population size prior to this point, from at least 50,000 BP to 5,000 BP followed by a sharp decline since then (Figure 3). The decline in effective female population size starts almost 4,000 years prior to the onset of whaling in the Cantabrian Sea.

**Figure 3.**
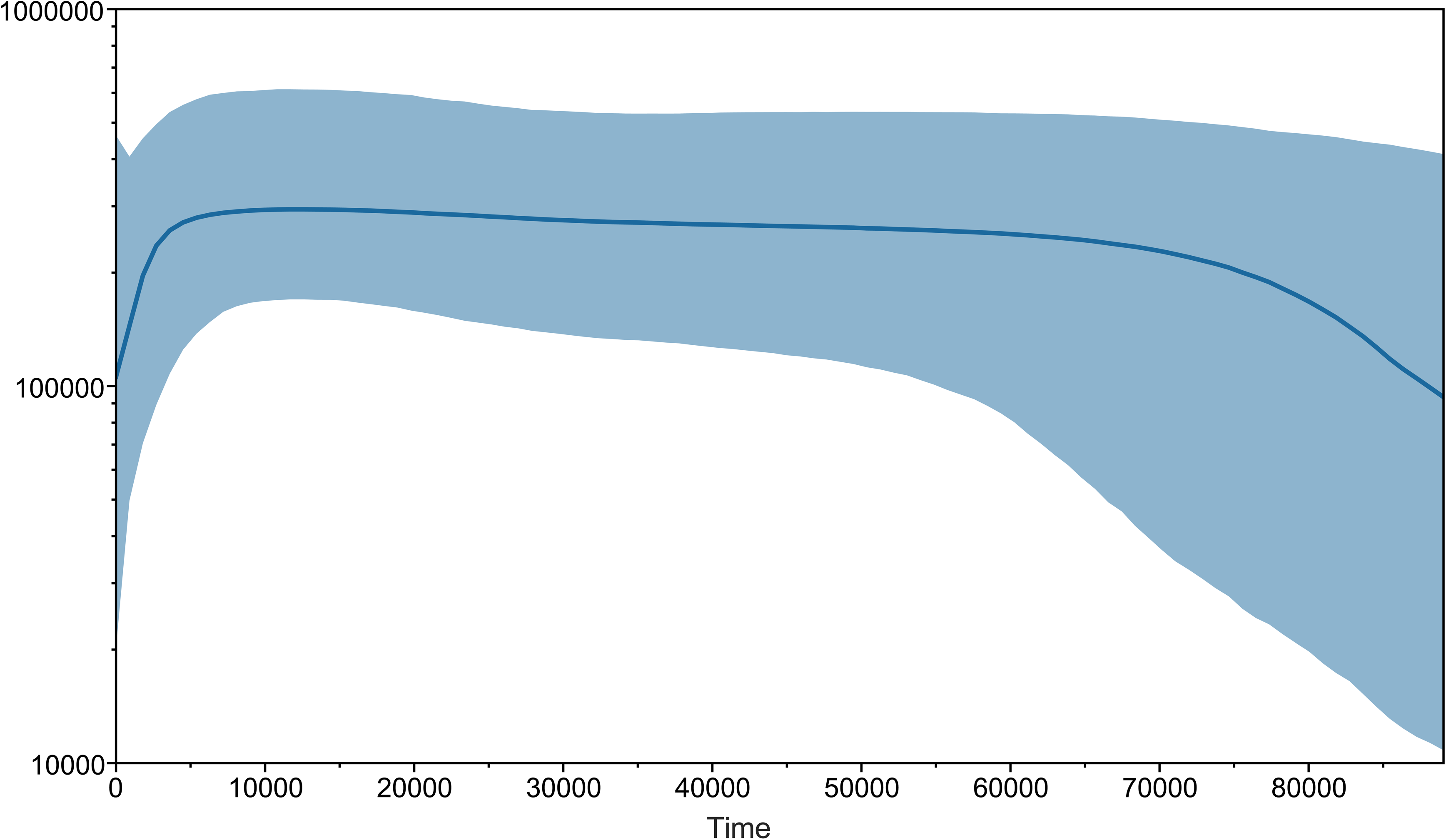
Bayesian skyline plot reconstructed from complete mitochondrial genomes of 41 North Atlantic Right whales. The dataset includes 24 modern individuals from the Northwest Atlantic and 17 historical specimens from the Cantabrian Sea in the Northeast Atlantic. The plot depicts changes in female effective population size (Ne) through time, showing a sharp decline in genetic diversity, beginning around 5000 years before present (BP).

**Figure 4.**
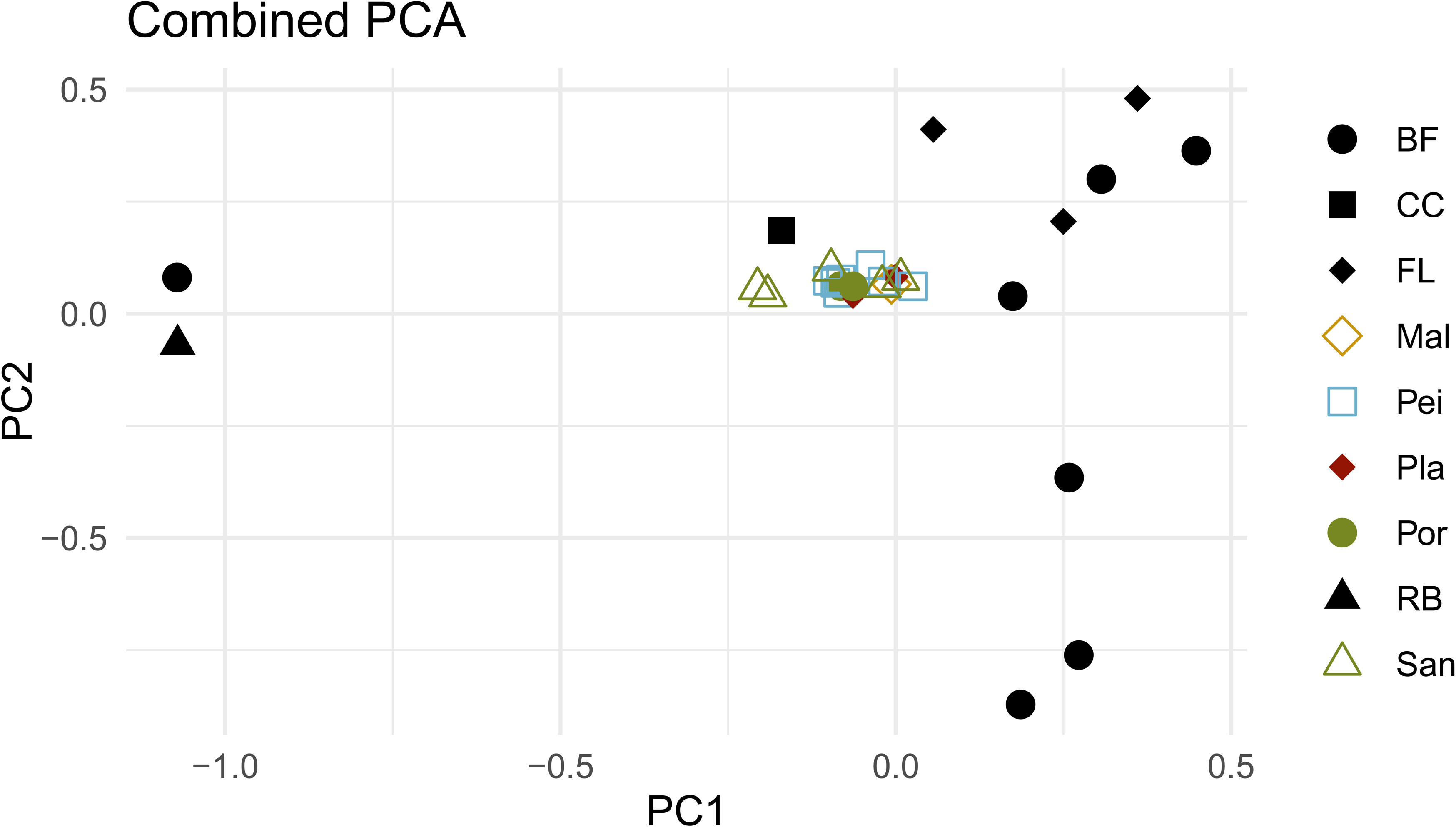
Procrustes-aligned principal component analysis (PCA) of genomic variation in 17 historical and 12 modern North Atlantic right whale specimens. The plot depicts genomic relationships among individuals after Procrustes alignment of PCA coordinates, highlighting similarities and differences between modern (feeding, bright blue; breeding, navy blue) and historical samples. Each point represents an individual positioned according to the first two principal components. The partial overlap between groups suggests continuity through time and the presence of a panmictic population.

Posterior distributions of demographic parameters indicate that the reduction in mitochondrial diversity is not the result of ancient demographic fluctuations but reflects a recent bottleneck. The near absence of previously common haplotypes in the modern population supports a scenario of lineage loss likely associated with the extirpation of local subpopulations, consistent with the intense whaling pressure of recent centuries. Therefore, our results provide genetic corroboration for historical accounts of intensive whaling and suggest that the demographic impacts of this exploitation are still evident in the genetic structure of the extant *Eubalaena glacialis* population.

Previous studies with more limited datasets that did not include historical samples had pointed to the potential negative impact of historical whaling on the NARW diversity. For example, a study examining six historical whale specimens from the late 19th century concluded that only a relatively modest change in maternal lineage diversity has occurred in North Atlantic right whales over the past century, and any substantial reduction in genetic variation took place prior to the late nineteenth century (Rosenbaum et al., 2000).

Furthermore, demographic reconstructions using both MSMC2 and STAIRWAY PLOT based on whole genome resequencing data from modern individuals confirm the reduction in effective population size in the past thousands of years as well as a continuous small effective population size for the species in great contrast to the southern right whale (Crossman et al., 2024) this decline to the onset and expansion of whaling on both sides of the Atlantic.

### Population structure

Population genetic parameters based on complete mitogenomes (n = 41) indicate greater genetic diversity among ancient individuals, with 17 haplotypes and 76 segregating sites, resulting in higher haplotype diversity (Hd = 1.00) and Watterson’s theta (Θ = 20.1) compared to the modern population (5 haplotypes, 29 segregating sites, Hd = 0.656, Θ = 11.9). Modern genetic diversity seems to be reduced when compared to Medieval samples. Even though the Northeast Atlantic population of NARW is now functionally extinct and the extant Northwest Atlantic population is quite small, part of the ancient diversity is still present today.

A Procrustes transformation of 17 individual PCAs obtained using the nuclear SNPs of 12 modern individuals and each of the 17 historical samples, reveals no structure between Northeastern and Northwestern NARW populations. This suggests that NARW is a panmictic species, and historically maintained by high levels of gene flow between eastern and western North Atlantic populations. Recently, a study by Crossman et al. (2024) based on whole genome resequencing of modern individuals also suggested that there is no population structure in this species and a previous study on one western Atlantic NARW from the 16^th^ century pointed to genetic continuity as all the alleles from 27 microsatellites analysed were found in present-day populations including with similar heterozygosity levels (McLeod et al., 2010).

### Sexing

Genetic sex determination identified seven individuals as male and ten as female, with no undetermined cases (Table S2). While historical accounts suggest a preferential targeting of females—particularly lactating individuals accompanied by calves (Aguilar, 1986; Harcourt et al., 2019) —the relatively high proportion of males (41%) in the sample suggests a more complex hunting dynamic. This pattern may reflect the incidental capture of male calves accompanying females, or alternatively, that adult males were also regularly hunted, either opportunistically or intentionally.

## Conclusion

Our genomic analysis of historical North Atlantic right whale specimens provides direct evidence of the species’ former panmictic structure and extensive gene flow across the Atlantic. The marked reduction in mitochondrial diversity and the near disappearance of common haplotypes in the modern population reveal severe lineage loss, consistent with the extirpation of regional subpopulations under centuries of intensive whaling. Demographic reconstructions indicate that this decline was not driven by ancient fluctuations but reflects a recent, human-induced bottleneck whose genetic signature persists today. By integrating ancient DNA with modern genomic data, we demonstrate that the extinction of the northeastern population and the erosion of diversity in the surviving western population are among the most profound anthropogenic impacts on any large marine vertebrate. These findings underscore the urgency of conservation measures for the critically endangered North Atlantic right whale and highlight the value of historical genomics in reconstructing population collapse and informing recovery strategies.

## Acknowledgements

This research was partially supported by the Strategic Funding UIDB/04423/2020 and UIDP/04423/2020, through national funds provided by FCT and European Regional Development Fund (ERDF), in the framework of the programme PT2020. PFC was partially supported by national funds through FCT, I.P., under the Scientific Employment Stimulus Initiative, reference CEECIND/01799/2017, 2023.05877.CEECIND and SARDINOMICS 2022.03142.PTDC. Sequencing data was obtained thanks to an EASI Genomics grant PID 14751, conceded to PFC. RDF thanks the VILLUM FONDEN for the Centre for Global Mountain Biodiversity grant no 25925.

---

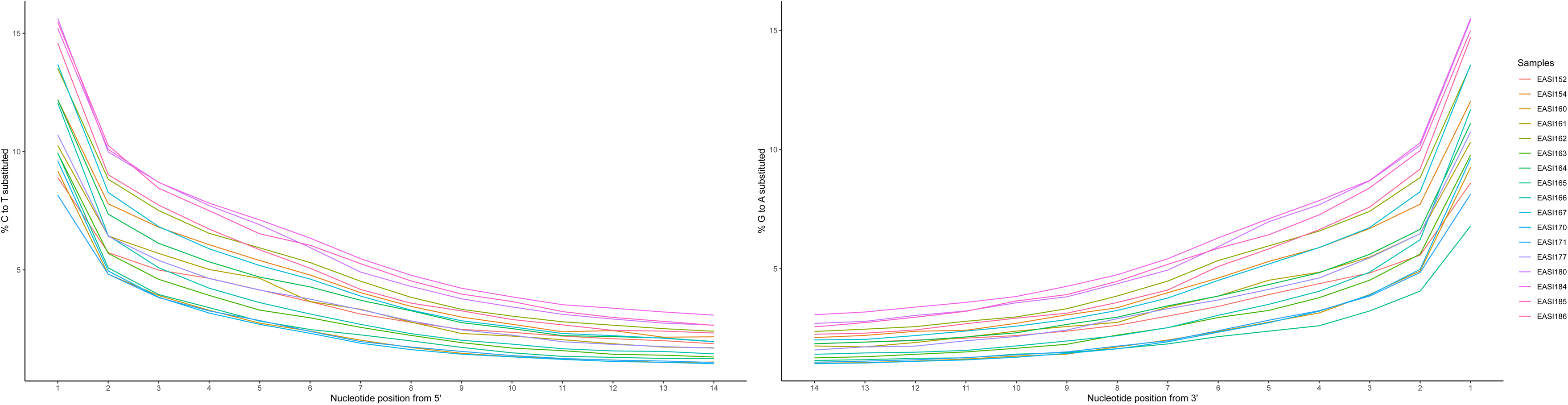

## References

Aguilar, A., 1986. A review of old Basque whaling and its effect on the right whales (Eubalaena glacialis) of the North Atlantic. Reports of the International Whaling Commitee, Special Issue 10, 191–199.

Bandelt, H.-J., Forster, P., Röhl, A., 1999. Median-joining networks for inferring intraspecific phylogenies. Mol Biol Evol 16, 37–48.

Béland, S.L., Frasier, B.A., Darling, J.D., Frasier, T.R., 2020. Using pre- and postexploitation samples to assess the impact of commercial whaling on the genetic characteristics of eastern North Pacific gray and humpback whales and to compare methods used to infer historic demography. Mar Mamm Sci 36, 398–420. 10.1111/mms.12652

Bouckaert, R., Vaughan, T.G., Barido-Sottani, J., Duchêne, S., Fourment, M., Gavryushkina, A., Heled, J., Jones, G., Kühnert, D., De Maio, N., Matschiner, M., Mendes, F.K., Müller, N.F., Ogilvie, H.A., du Plessis, L., Popinga, A., Rambaut, A., Rasmussen, D., Siveroni, I., Suchard, M.A., Wu, C.-H., Xie, D., Zhang, C., Stadler, T., Drummond, A.J., 2019. BEAST 2.5: An advanced software platform for Bayesian evolutionary analysis. PLoS Comput Biol 15, e1006650. 10.1371/journal.pcbi.1006650

Ciriquiain-Gaiztarro, M., 1961. Los vascos en la pesca de la ballena. Biblioteca Vascongada de los Amigos del País.

Crossman, C.A., Fontaine, M.C., Frasier, T.R., 2024. A comparison of genomic diversity and demographic history of the North Atlantic and Southwest Atlantic southern right whales. Mol Ecol 33, e17099. 10.1111/MEC.17099

Drummond, A.J., Rambaut, A., Shapiro, B., Pybus, O.G., 2005. Bayesian coalescent inference of past population dynamics from molecular sequences. Mol Biol Evol 22, 1185–1192.

Fellows Yates, J.A., Lamnidis, T.C., Borry, M., Andrades Valtueña, A., Fagernäs, Z., Clayton, S., Garcia, M.U., Neukamm, J., Peltzer, A., 2021. Reproducible, portable, and efficient ancient genome reconstruction with nf-core/eager. PeerJ 9, e10947. 10.7717/peerj.10947

Frankham, Richard., Ballou, J.D.., Briscoe, D.A.., 2010. Introduction to conservation genetics 618.

Frasier, B.A., Springate, L., Frasier, T.R., Brewington, S., Carruthers, M., Edvardsson, R., Harrison, R., Kitchener, A.C., Mainland, I., Szabo, V.E., 2022. Genetic examination of historical North Atlantic right whale (*Eubalaena glacialis*) bone specimens from the eastern North Atlantic: Insights into species history, transoceanic population structure, and genetic diversity. Mar Mamm Sci 38, 1050–1069. 10.1111/mms.12916

Freitas, C., Skogen, M.D., Sigurðsson, G.M., Biuw, M., Haug, T., Lindblom, L., Gundersen, K., 2025. Impact of baleen whales on ocean primary production across space and time. Proc Natl Acad Sci U S A 122, e2505563122. 10.1073/PNAS.2505563122;WGROUP:STRING:PUBLICATION

Harcourt, R., van der Hoop, J., Kraus, S., Carroll, E.L., 2019. Future directions in Eubalaena spp.: Comparative research to inform conservation. Front Mar Sci 6, 417957. 10.3389/FMARS.2018.00530/FULL

Korneliussen, T.S., Albrechtsen, A., Nielsen, R., 2014. ANGSD: analysis of next generation sequencing data. BMC Bioinformatics 15, 356.

Leigh, J.W., Bryant, D., 2015. popart: full-feature software for haplotype network construction. Methods Ecol Evol 6, 1110–1116.

Li, H., Durbin, R., 2009. Fast and accurate short read alignment with Burrows–Wheeler transform. Bioinformatics 25, 1754–1760.

López, A., 2014. Historia ambiental antigua das baleas do Atlántico Norte. Eubalaena 14, 3–27.

Malik, S., Brown, M.W., Kraus, S.D., White, B.N., 2000. Analysis of mitochondrial DNA diversity within and between North and South Atlantic right whales. Mar Mamm Sci 16, 545–558. 10.1111/J.1748-7692.2000.TB00950.X

Maricic, T., Whitten, M., Pääbo, S., 2010. Multiplexed DNA sequence capture of mitochondrial genomes using PCR products. PLoS One 5, e14004.

McCartney, A.P., 1980. The nature of Thule Eskimo whale use. Arctic 517–541.

McCartney, A.P., Savelle, J.M., 1993. Bowhead whale bones and Thule Eskimo subsistence–settlement patterns in the central Canadian Arctic. Polar Record 29, 1–12.

McLeod, B.A., Brown, M.W., Frasier, T.R., White, B.N., 2010. DNA profile of a sixteenth century western North Atlantic right whale (Eubalaena glacialis). Conservation Genetics 11, 339–345. 10.1007/s10592-009-9811-6

Meisner, J., Albrechtsen, A., 2018. Inferring Population Structure and Admixture Proportions in Low-Depth NGS Data. Genetics 210, 719–731. 10.1534/genetics.118.301336

Meyer, M., Kircher, M., 2010. Illumina sequencing library preparation for highly multiplexed target capture and sequencing. Cold Spring Harb Protoc 2010, pdb. prot5448.

Neukamm, J., Peltzer, A., Nieselt, K., 2021. DamageProfiler: fast damage pattern calculation for ancient DNA. Bioinformatics 37, 3652–3653. 10.1093/bioinformatics/btab190

Pettis, H., Pace, R.I., Hamilton, P., 2022. North Atlantic Right Whale Consortium 2021 Annual Report Card.

Rambaut, A., Drummond, A.J., Xie, D., Baele, G., Suchard, M.A., 2018. Posterior summarization in Bayesian phylogenetics using Tracer 1.7. Syst Biol 67, 901–904. 10.1093/sysbio/syy032

Rastogi, T., Brown, M.W., McLeod, B.A., Frasier, T.R., Grenier, R., Cumbaa, S.L., Nadarajah, J., White, B.N., 2004. Genetic analysis of 16th-century whale bones prompts a revision of the impact of Basque whaling on right and bowhead whales in the western North Atlantic. Can J Zool 82, 1647–1654.

Reeves, R.R., Smith, T.D., 2007. A Taxonomy of World Whaling, in: Estes, J.A., DeMaster, D.P., Doak, D.F. (Eds.), Whales, Whaling, and Ocean Ecosystems. University of California Press, Berkeley, pp. 82–101. 10.1525/california/9780520248847.003.0008

Rey-Iglesia, A., Martínez-Cedeira, J., López, A., Fernández, R., Campos, P.F., 2018. The genetic history of whaling in the Cantabrian Sea during the 13th–18th centuries: Were North Atlantic right whales (*Eubalaena glacialis*) the main target species? J Archaeol Sci Rep 18, 393–398. 10.1016/j.jasrep.2018.01.034

Roman, J., McCarthy, J.J., 2010. The Whale Pump: Marine Mammals Enhance Primary Productivity in a Coastal Basin. PLoS One 5, e13255. 10.1371/journal.pone.0013255

Rosenbaum, H.C., Egan, M.G., Clapham, P.J., Brownell, R.L., Malik, S., Brown, M.W., White, B.N., Walsh, P., Desalle, R., 2000. Utility of North Atlantic Right Whale Museum Specimens for Assessing Changes in Genetic Diversity. Conservation Biology 14, 1837–1842. 10.1111/J.1523-1739.2000.99310.X;CTYPE:STRING:JOURNAL

Sasaki, T., Nikaido, M., Hamilton, H., Goto, M., Kato, H., Kanda, N., Pastene, L.A., Cao, Y., Fordyce, R.E., Hasegawa, M., 2005. Mitochondrial phylogenetics and evolution of mysticete whales. Syst Biol 54, 77–90.

Schubert, M., Lindgreen, S., Orlando, L., 2016. AdapterRemoval v2: rapid adapter trimming, identification, and read merging. BMC Res Notes 9, 88. 10.1186/s13104-016-1900-2

Seersholm, F.V., Pedersen, M.W., Søe, M.J., Shokry, H., Mak, S.S.T., Ruter, A., Raghavan, M., Fitzhugh, W., Kjær, K.H., Willerslev, E., Meldgaard, M., Kapel, C.M.O., Hansen, A.J., 2016. DNA evidence of bowhead whale exploitation by Greenlandic Paleo-Inuit 4,000 years ago. Nat Commun 7, 1–9. 10.1038/ncomms13389

Skovrind, M., Louis, M., Ferguson, S.H., Glazov, D.M., Litovka, D.I., Loseto, L., Meschersky, I.G., Miller, M.M., Petr, M., Postma, L., Rozhnov, V. V., Scott, M., Westbury, M. V., Szpak, P., Friesen, T.M., Lorenzen, E.D., 2024. Elucidating the sustainability of 700 y of Inuvialuit beluga whale hunting in the Mackenzie River Delta, Northwest Territories, Canada. Proceedings of the National Academy of Sciences 121. 10.1073/pnas.2405993121

Solazzo, C., Fitzhugh, W., Kaplan, S., Potter, C., Dyer, J.M., 2017. Molecular markers in keratins from Mysticeti whales for species identification of baleen in museum and archaeological collections. PLoS One 12, e0183053. 10.1371/journal.pone.0183053

Speller, C., van den Hurk, Y., Charpentier, A., Rodrigues, A., Gardeisen, A., Wilkens, B., McGrath, K., Rowsell, K., Spindler, L., Collins, M., Hofreiter, M., 2016. Barcoding the largest animals on Earth: ongoing challenges and molecular solutions in the taxonomic identification of ancient cetaceans. Philosophical Transactions of the Royal Society B: Biological Sciences 371, 20150332. 10.1098/rstb.2015.0332

Sueyro, N., Crespo, E.A., Arias, M., Coscarella, M.A., 2018. Density-dependent changes in the distribution of Southern Right Whales (*Eubalaena australis*) in the breeding ground Peninsula Valdés. PeerJ 6, e5957. 10.7717/peerj.5957

Valdés Hansen, F., 2010. Los balleneros en Galicia (siglos XIII al XX). Colleción Galicia Histórica. Fundación Pedro Barrié de la Maza, A Coruña.

Waldick, R.C., Kraus, S., Brown, M., White, B.N., 2002. Evaluating the effects of historic bottleneck events: an assessment of microsatellite variability in the endangered, North Atlantic right whale. Mol Ecol 11, 2241–2249. 10.1046/J.1365-294X.2002.01605.X

