## Supplementary Figure 2 for "Historic Overexploitation, Genetic Erosion and Local extinction: Palaeogenomic Insights into the Decline of *Eubalaena glacialis* in the Northeast Atlantic"

Table S2 – Depth of coverage ratio between sex chromosome X and the autosome chromosome 7. A ratio below 0.6 indicates a male, while a ratio above 0.8 indicates a female.

| Sample | NC_083736 (X) coverage | NC_083722 (chr7) coverage | ratio |
| --- | --- | --- | --- |
| EASI152 | 0.049497538 | 0.057432787 | 0.861834 |
| EASI 154 | 0.034240484 | 0.03847125 | 0.890028 |
| EASI 160 | 0.050012382 | 0.0541732 | 0.923194 |
| EASI 161 | 0.009416349 | 0.016236107 | 0.579963 |
| EASI 162 | 0.050399609 | 0.099583229 | 0.506105 |
| EASI 163 | 0.039442961 | 0.075715107 | 0.520939 |
| EASI 164 | 0.065766794 | 0.076813008 | 0.856193 |
| EASI 165 | 0.069425625 | 0.07407116 | 0.937283 |
| EASI 166 | 0.115800844 | 0.13127326 | 0.882136 |
| EASI 167 | 0.064233135 | 0.069696124 | 0.921617 |
| EASI 170 | 0.0559407 | 0.105939252 | 0.528045 |
| EASI 171 | 0.101066768 | 0.190411264 | 0.530781 |
| EASI 177 | 0.033003476 | 0.038122831 | 0.865714 |
| EASI 180 | 0.034952079 | 0.039584897 | 0.882965 |
| EASI 184 | 0.131643396 | 0.149575699 | 0.880112 |
| EASI 185 | 0.012220264 | 0.023902259 | 0.51126 |
| EASI 186 | 0.01253724 | 0.023248365 | 0.539274 |
